# Reaction Conditions Promoting the Specific Detection of SARS-CoV-2 NendoU Enzymatic Activity

**DOI:** 10.1101/2022.05.06.490915

**Authors:** Nodar Makharashvili, James O. McNamara

## Abstract

Methods that enable rapid detection of SARS-CoV-2 provide valuable tools for detecting and controlling Covid-19 outbreaks and also facilitate more effective treatment of infected individuals. The predominant approaches developed use PCR to detect viral nucleic acids or immunoassays to detect viral proteins. Each approach has distinct advantages and disadvantages, but alternatives that do not share the same limitations could enable substantial improvements in outbreak detection and management. For instance, methods that have comparable sensitivity to PCR, but that are not prone to the false-positive results that stem from the tendency of PCR to detect molecular degradation products could improve accurate identification of infected individuals. An alternative approach with potential to achieve this entails harnessing the unique enzymatic properties of SARS-CoV-2 enzymes to generate SARS-Cov-2-specific signals that indicate the presence of the virus. This route benefits from the high sensitivity provided by enzymatic signal amplification and also the fact that signal is generated only by intact viral enzymes, not degradation products. Here, we demonstrate enzymatic reaction conditions that enable the preferential detection of NendoU of SARS-CoV-2, versus several of its orthologues, with a fluorogenic oligonucleotide substrate. These compositions provide a possible technical foundation for a novel approach for detecting SARS-CoV-2 that has distinct advantages from current approaches.

## Introduction

Rapid and inexpensive assays that detect the SARS-CoV-2 coronavirus in human samples are valuable tools that enable identification of infected individuals and monitoring of infection prevalence in communities (Peeling et al., 2022). Detection of infections enables the selective isolation of infected subjects from uninfected individuals which can greatly reduce disease transmission. Efficient infection detection can also enable treatment early in the course of illness, improving treatment outcomes. Moreover, viral detection methods can be used to monitor the prevalence of infections in communities, providing information that can be used to guide behavioral adjustments at the community level to limit the impact of outbreaks.

Several types of detection assays have been developed for SARS-CoV-2, including PCR-based assays that detect viral RNA, antigen assays that detect SARS-CoV-2 proteins, usually proteins of the viral capsid, and assays that detect SARS-CoV-2 infection signatures in the cells and molecules of the human immune system (i.e., host-response tests such as antibody tests) (Peeling et al., 2022). The PCR and antigen tests are the assay types that are most useful for identifying individuals with active infections. PCR tests are more sensitive and are considered the gold-standard tests. However, PCR tests are known to produce false-positive results based on their detection of nucleic acid degradation products (i.e., inactive virus), resulting in the unnecessary and often costly isolation of individuals who do not pose a viral transmission risk.

## Results

This study describes assay conditions that enable the specific detection of SARS-CoV-2 NendoU (Bhardwaj et al., 2008; Bhardwaj et al., 2006; Ivanov et al., 2004) via its digestion of a fluorogenic poly-uridine (RNA) oligonucleotide substrate. The conditions are the product of our evaluations of various oligonucleotide substrates and buffer conditions that were tested to identify compositions that selectively promote the enzymatic activity of SARS-CoV-2 NendoU versus the activities of its orthologues of distinct human coronaviruses. To develop methods for specific detection of SARS-CoV-2 NendoU, recombinant NendoU of SARS-CoV-2 was prepared together with recombinant versions of the related NendoU orthologues of the four human coronaviruses that are in common circulation: 229E, NL63, OC43 and HKU1. Comparison of the amino acid sequences of these NendoU orthologues to that of the SARS-CoV-2 version indicate identities of 39% to 45%. Two reaction buffers that enable preferential detection of the SARS-CoV-2 NendoU orthologue were identified (**Figure 1**). The background-subtracted activity of SARS-CoV-2 NendoU exceeded that of its orthologues by at least 3-4 fold with each of the buffers. Both buffers include divalent cations that can serve as inhibitors of off-target nucleases (Yang, 2011) but permit the activity of NendoU of SARS-CoV-2. Additionally, the NendoU activity occurred in the presence of the disulfide-bond reducing reagents, beta-mercaptoethanol (MnCl2 buffer) or TCEP (NiCl2 buffer). This is notable because some off-target nucleases, such as RNase A, require disulfide bonds to exhibit enzymatic activity (Yang, 2011). Inclusion of disulfide bond reducing reagents may therefore be used to promote assay specificity for SARS-CoV-2 NendoU.

**Figure 1.**
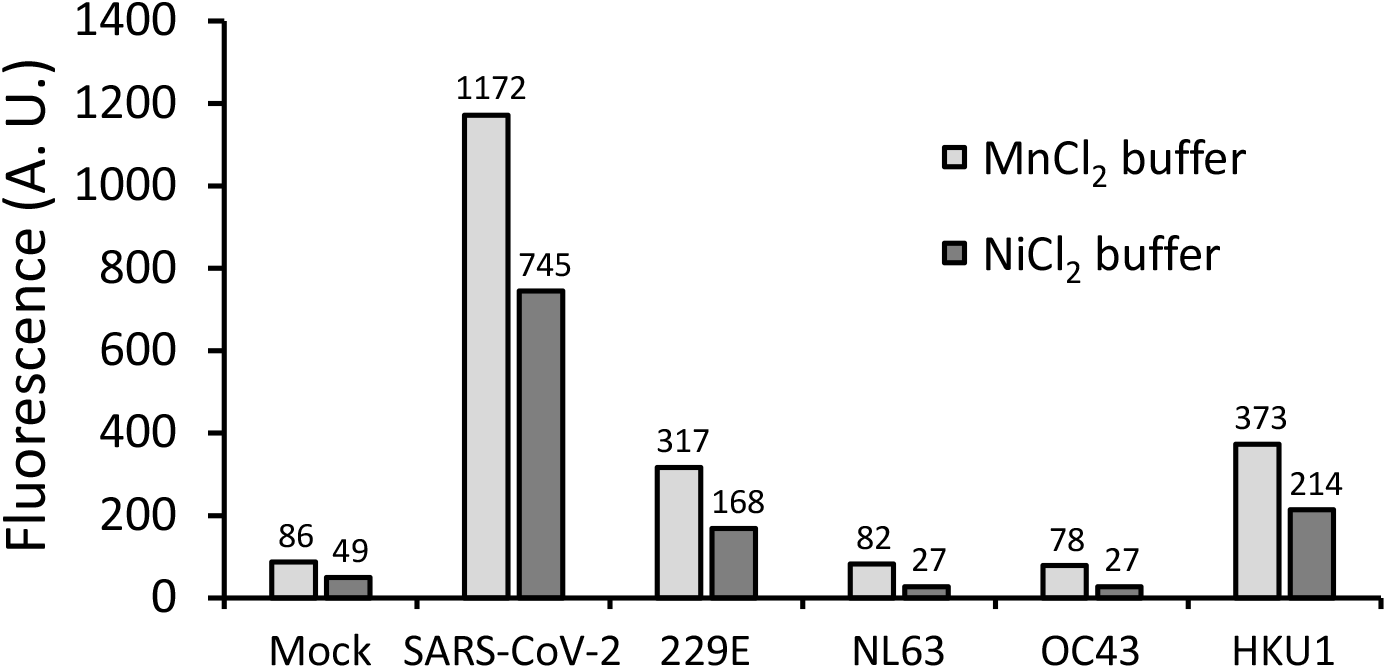
Nuclease reaction buffers promote detection specificity for SARS-CoV-2 NendoU versus its human coronavirus orthologues. 5 μL of 1 μM of each of the indicated NendoU orthologues was combined with 45 μL of either a MnCl_2_ buffer (50 mM Tris pH = 7.0, 20 mM MnCl_2_, 250 mM KCl, 20 mM beta-mercaptoethanol) or a NiCl_2_ buffer (50 mM Tris pH = 7.0, 40 mM NiCl_2_, 250 mM KCl, 5 mM TCEP, 10 mM PVSA). Reactions also contained 0.1 μM PolyU fluorogenic RNA probe. Fluorescence was measured after incubation for 30 minutes at 37 °C.

## Discussion

The enzymatic reaction conditions demonstrated here enable the preferential detection of the SARS-CoV-2 NendoU nuclease versus its orthologues that are produced by distinct human coronaviruses with fluorogenic oligonucleotide probes. This work may thereby provide the basis for a novel means of specifically detecting SARS-CoV-2 with key advantages over existing methods. Like antigen tests, assays that would detect SARS-CoV-2 via NendoU would detect the virus via the presence of a SARS-CoV-2 protein product. However, while antigen tests use antibodies to specifically bind and detect their protein target, a NendoU test would use a fluorogenic enzymatic substrate to specifically detect the NendoU protein via its unique enzymatic properties. Rather than using assay reagents such as secondary antibodies coupled to reporter molecules for detection, this nuclease assay approach harnesses the enzymatic activity of the target protein to generate detectable signal directly (Burghardt et al., 2016; Flenker et al., 2017; Hernandez et al., 2014; Lopez-Alvarez et al., 2020). The approach thus requires fewer molecular components, with the primary component consisting of an oligonucleotide substrate that is produced with chemical synthesis. Furthermore, nuclease assays require a functional enzymatic target to generate signal, reducing or eliminating the risk that false positive results elicited by degradation products of the target molecule will occur. While antigen tests have exhibited lower sensitivity than PCR, they have the potential to provide superior sensitivity because proteins (the molecular targets of antigen tests) are roughly ~2,000-fold more abundant than nucleic acids) (Marguerat et al., 2012).

Future work is needed to translate this approach into practice. Key objectives include validation in human sample matrices such as saliva and addressing any background probe digestion that may result from human-derived nucleases. As the oligonucleotide portion of the substrate consists of unmodified RNA, some means of preventing digestion by human RNases may be needed. One avenue for achieving this may be to introduce a NendoU-capturing step prior to the enzymatic reaction step. This could enable removal of off-target nucleases prior to the reaction step and has potential to substantially improve both specificity and sensitivity. This capture/measure approach has previously shown promise in a rapid assay for bacterial bloodstream infections (Burghardt et al., 2016).

## Materials and Methods

Plasmid constructs. Amino acid sequences of the NendoU (Nsp15) orthologues of the following human coronaviruses were obtained from the Uniprot.org database: SARS-CoV2; HCoV-229E; HCoV-NL63; HCoV-OC43; HCoV-HKU1. Protein accession numbers and amino acid numbers used (in brackets) are as follows: SARS-CoV2 (accession number: P0DTD1[6453 - 6798]); HCoV-229E (accession number: P0C6X1[6111 - 6458]); HCoV-NL63 (accession number: P0C6X5[6086 - 6429]); HCoV-OC43 (accession number: P0C6X6, [6422 - 6796]); HCoV-HKU1 (accession number: P0C6X3, [6480 - 6853]). DNA sequences encoding each protein were codon-optimized for bacterial expression, synthesized, and cloned into the pET28a expression plasmid as custom orders by GenScript Biotech.

Protein expression. The protein expression plasmids were transformed into BL21-pLys expression cells according to the manufacturer’s recommendations (Thermo Scientific). Transformed bacterial clones were expanded in LB supplemented with kanamycin at 37 °C, with shaking. After an initial outgrowth period, expression was induced with addition of Isopropyl-β-D-thiogalactopyranoside (IPTG) (Fisher Scientific) and the cultures were grown for an additional 20 hours at 18 °C.

Protein purification.The His-tagged nucleases were purified with standard methods. Briefly, the cells were harvested by centrifugation, lysed in the presence of protease inhibitors, the lysate was clarified with centrifugation and the proteins were captured with Ni-NTA agarose (Invitrogen). The agarose was washed and then proteins were eluted with imidazole-containing buffer. Protein products were analyzed with polyacrylamide gel electrophoresis (PAGE). Purified proteins were flash frozen in liquid nitrogen and stored at −80 °C.

Reaction conditions. Nuclease reactions were prepared by adding 5 μL of 1 μM of the indicated NendoU orthologues to 45 μL of either a MnCl2 buffer (50 mM Tris pH = 7.0, 20 mM MnCl2, 250 mM KCl, 20 mM beta-mercaptoethanol) or a NiCl2 buffer (50 mM Tris pH = 7.0, 40 mM NiCl2, 250 mM KCl, 5 mM TCEP, 10 mM PVSA). Reactions also contained PolyU fluorogenic RNA probe (11mer poly-uridine flanked with fluorophore and quencher; 0.1 μM concentration). Fluorescence was measured with a Synergy H1 Biotek plate-reader after incubation for 30 minutes at 37 °C. Fluorescence measures were acquired with standard green channel fluorescence wavelengths.

Reagents. NiCl2 was acquired from Alfa Aesar; KCl was acquired from Invitrogen; PVSA (poly(vinylsulfonic acid) was acquired from Polysciences. The following reagents were acquired from Fisher Scientific: 1 M Tris-HCl pH 7.0 stock solution, TCEP-HCl, MnCl2, beta-mercaptoethanol, NaCl.

The PolyU probe was acquired as a custom oligo from IDT (IDT) of Coralville, IA. This is an 11-mer RNA oligo flanked with a fluorophore and quenchers (5′- FAM-rUrUrUrUrUrUrUrUrUrUrU-IAbRQSp-3′, where FAM is fluorescein amidite, rU is a uridine nucleotide (RNA), and IAbRQSp is the Iowa Black RQ quencher connected with a spacer).

## Notes

### Competing Interest Statement

J.O.M. is the founder and CEO of Nuclease Probe Technologies (NPT), a biotechnology company developing rapid diagnostic assays for infectious diseases. N.M. is the Director of Research and Development at NPT.

